# Scaling down the microbial loop: data-driven modelling of growth interactions in a diatom-bacterium co-culture

**DOI:** 10.1101/2021.03.17.435777

**Authors:** Giulia Daly, Elena Perrin, Carlo Viti, Marco Fondi, Alessandra Adessi

**Affiliations:** Department of Agriculture, Food, Environment and Forestry, University of Florence, Piazzale delle Cascine 18, Florence, Italy; Department of Biology, University of Florence, Via Madonna del Piano 6, Sesto F.no, Florence, Italy

## Abstract

An intricate set of interactions characterizes marine ecosystems. One of the most important is represented by the so-called microbial loop, which includes the exchange of dissolved organic matter (DOM) from phototrophic organisms to heterotrophic bacteria. Here, it can be used as the major carbon and energy source. Arguably, this interaction is one of the foundations of the entire ocean food-web. Carbon fixed by phytoplankton can be redirected to bacterial cells in two main ways; either i) bacteria feed on dead (eventually lysed) phytoplankton cells or ii) DOM is actively released by phytoplankton cells (a widespread process that may result in up to 50% of the fixed carbon leaving the cell). In this work, we have set up a co-culture of the model diatom *Phaeodactylum tricornutum* and the model chemoheterotrophic bacterium *Pseudoalteromonas haloplanktis* TAC125 and used this system to study the interactions between these two representatives of the microbial loop. We show that the bacterium can indeed thrive on diatom-derived carbon and that this growth can be sustained by both diatom dead cells and diatom-released compounds. These observations were formalized in a network of putative interactions between *P. tricornutum* and *P. haloplanktis* and implemented in a mathematical model that reproduces the observed co-culture dynamics, suggesting that our hypotheses on the interactions occurring in this two-player system can accurately explain the experimental data.

## Introduction

The complex network of nutrients exchange among marine organisms is usually referred to as the microbial loop (Azam et al. 1983; Fenchel 2008). The foundation of this intricate system is represented by microbial phototroph–heterotroph interactions that permit the biogeochemical cycling of elements and, more in general, represent a model for understanding and predicting ocean processes on a global scale (Christie-Oleza et al. 2017). In general, bacteria thrive on algal-excreted organic carbon and, in return, the algae may benefit from bacteria-recycled nutrients and other metabolites (Amin, Parker, and Armbrust 2012; Bell and Mitchell 1972). Indeed, phytoplankton represents one of the main sources of carbon for bacteria in the ocean thanks to the carbon they may provide to the overall dissolved organic matter (DOM) pool through exudation or death-related processes (Azam et al. 1983; Landa et al. 2017). In natural settings, these latter processes may be represented by predators feeding (Moller 2004), cell lysis (Bettarel et al. 2005; Bratbak, Egge, and Heldal 1993; Gobler et al. 1997), and cell death in general (Veldhuis, Kraay, and Timmermans 2001). Moreover, living phytoplankton cells exude a significant proportion, and under some circumstances the majority, of their photosynthate into the surrounding environment (Fogg, 1983; Wood & Van Valen, 1990). It has been estimated that a large fraction (up to 50%) of the fixed carbon is actually released in the surrounding environment and thus could be used by heterotrophic bacteria. However, the exact amount, composition and release rates of phytoplankton-derived DOM are far from being completely understood. A plethora of experimental and computational approaches have been used to develop models accounting for the release rates of DOM by phytoplankton (Biddanda and Benner 1997; Flynn, Clark, and Xue 2008), for the composition of DOM (Omta et al. 2020), for the interaction with surrounding heterotrophs (Fondi and Di Patti 2019; Moejes et al. 2017; Morán et al. 2001). Although all these works converge on some aspects, such as the emergence of metabolic interdependencies and higher-order interactions (Croft et al. 2005; Mickalide and Kuehn 2019), some points remain still debated. These include, for example, the contribution of phytoplankton cell lysis to the overall DOM pool, the connection between the physiological state of the cell and the amount of carbon released, the role of the heterotrophic community in the shaping of this nutrient loop.

Among the others, two important groups of marine microbes are known to interact and play a major role in defining the microbial loop dynamics, i.e. bacteria and diatoms (Amin, Parker, and Armbrust 2012). Diatoms and bacteria have been shown to possess many possible ways of interactions, including synergistic (Croft et al. 2005), parasitic (Paul and Pohnert 2011) and competitive (Bratbak and Thingstad 1985) ones. Disentangling their network of interactions and elucidating how each of these two players influences the physiology of the other is key for deciphering oceanic nutrient fluxes and biogeochemical cycles. Also, the composition of the diatom-associated bacterial community has been shown to play a role in regulating the physiological status of this biological system (Behringer et al. 2018; Moejes et al. 2017; Rooney-Varga et al. 2005) and to be the subject of specific regulatory rules. In a recent work, Shibl et al. (2020) showed that diatom exudates might tune microbial communities and select specific bacteria of their associated consortium. This is achieved through the secretion of secondary metabolites that promote the proliferation of selected bacteria and demote others (Shibl et al. 2020). Further, Moejes et al. (2017) have characterized the microbiome associated with the model pennate diatom *Phaeodactylum tricornutum* and proposed a network of putative interactions between the diatom and the main bacterial taxa found in the community. Then, this was formalized in a mathematical model able to qualitatively reproduce the observed community dynamics, also through the inference of nutrients exchange among the representatives of this biological association. Despite providing valuable hints on the possible, high-level interactions among the different community members, the complexity of these systems makes it hard to specifically address the interactions occurring between the diatom and specific bacterial representatives. In the presence of a co-culture that embeds a complex bacterial community, for example, it is not possible to understand which microbial representative(s) is actually feeding on diatom-released DOM or sequestering specific micronutrients from the system.

To overcome these limitations, we have set up a co-culture of two model representatives of the microbial loop, the diatom *P. tricornutum* and the chemoheterotrophic bacterium *Pseudoalteromonas haloplanktis* TAC125. Importantly, this bacterium has been used as a model system to study the interaction between nutrients and bacteria in the marine environment (Perrin et al. 2020; Stocker et al. 2008). In a previous work (Fondi and Di Patti 2019), we have simulated the putative metabolic cross-talks of this phototroph-heterotroph system using constraint-based metabolic modelling and showed that this combined metabolic reconstruction was able to suggest coarse-grained interactions of this simplified microbial community. Here, we build on this previous knowledge and experimentally test the capability of these two microbes to coexist in the same co-culture and shed light on the trophic interactions among them. We demonstrate that the bacterium can indeed thrive on diatom-derived carbon and that this growth can be sustained by both diatom dead cells and diatom-released compounds. On the contrary, the bacterium seems not to influence the growth of the diatom in the tested co-culture conditions. These observations were formalized in a network of putative interactions between *P. tricornutum* and *P. haloplanktis* that, in turn, was implemented as a mathematical model reproducing the observed co-culture dynamics. We aimed to test if our hypotheses on the interactions occurring in this two-player system could explain the experimental data.

## Results and Discussion

### Growth dynamics of *P. haloplanktis* and *P. tricornutum* co-culture

In this work we have set up a two-members co-culture system, composed of the diatom *P. tricornutum* and the chemoheterotrophic bacterium *P. haloplanktis* TAC125 for 28 days with no additional carbon sources, ensuring bacterial growth was dependent upon diatom released organic molecules. The co-culture was regularly sampled every 7 days (from day 0 to day 28, 5 time points). In order to study the interactions of *P. tricornutum* and *P. haloplanktis*, it was necessary to establish the optimal growth conditions to co-culture these two microorganisms, including medium composition, pH, temperature, and nutritional dependency.

The population dynamics of *P. tricornutum* and *P. haloplanktis* in co-culture and as single cultures were obtained. During the first phase of the co-culture (day 0 to 14), the bacterium *P. haloplanktis* showed a steady number of cells, with no evident growth, followed by a remarkable increase from day 14 to day 28, with respect to the bacterial negative control (Figure 1A). As a matter of fact, we found a statistically significant difference in the number of *P. haloplanktis* cells when comparing the bacterium-diatom co-culture and the *P. haloplanktis* negative control (t-test, p-value = 0,0024 and p-value = 0,0007 for 21 and 28 days, respectively). The bacterial positive control reached a cell density of 6,7 × 10^7^ CFU/ml by day 14, after which it started to decrease until the end of the cultivation (Supplementary Table S1).

**Figure 1:**
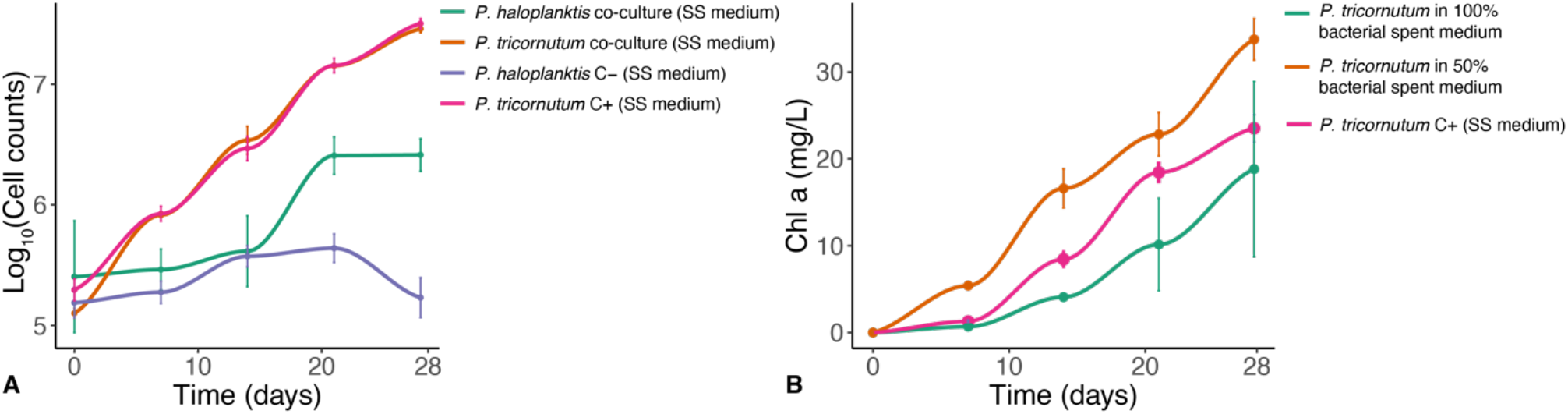
A) Co-culture data (including controls) of *P. tricornutum* and *P. haloplanktis* growth. Co-culture, *P. tricornutum* positive control and *P. haloplanktis* negative control were all cultured in SS medium (with no carbon source added). Error bars, standard deviation of triplicate cultures. B) Chlorophyll a content in *P. tricornutum* grown in non-diluted bacterial spent medium (100%) and 50% diluted medium. *P. tricornutum* positive control grown in fresh SS medium. Error bars, standard deviation of duplicate cultures.

*P. tricornutum* growth was unaffected when co-cultured with *P*.*haloplanktis* for the entire experiment. Overall, *P. tricornutum* cell number (Figure 1A) and other features that can be used for monitoring algal growth, such as the concentration of chlorophyll *a*, which is a proxy for phytoplankton abundance (Cole, Findlay, and Pace 1988)(Supplementary Figure S1), and the culture pH, were not affected by the presence of *P. haloplanktis* cells. In our model co-culture, the presence of the bacterium did not seemingly influence the diatom growth, not even in the last stages, after the bacterial increased growth. One possible explanation for these observations is that the bacteria in the first part of the cultivation were able to survive in co-culture with diatoms, maintaining a stable cell density (likely using small amounts of diatom-derived organic carbon present in the medium), later, in the second phase (days 14-28), when the diatoms have increased cell number and have released a sufficient DOM to sustain the bacterial growth, the bacteria actively started to grow. These findings suggest that the carbon sources required by *P. haloplaktis* for its growth were mainly provided during the late phase of the experiment (first part of the diatom stationary phase), when generally several inorganic nutrients become limited and phytoplankton release increasing amounts of DOM, used as carbon and energy source by heterotrophic bacteria (Bratbak and Thingstad 1985; Diner et al. 2016). Previous studies on *P. tricornutum* have confirmed that the concentration of organic carbon released by the diatom increases after the cells get into the stationary phase (Chen and Wangersky 1996); (Pujo-Pay M, Conan P, and Raimbault P 1997). Earlier co-culturing studies have shown that dynamics between phytoplankton and bacteria have occurred late in the growth cycles (Grossart and Simon 2007; Wang et al. 2014). For example, after a first mutualistic phase, Wang *et al*. (2014) highlighted a second pathogenic phase (day 21-36) which broke the balance existing between the bacterium and the dinoflagellate.

Within phytoplankton-bacteria interactions, bacteria can provide the phytoplankton cells with limiting macronutrients via remineralization (Legendre and Rassoulzadegan 1995) as well as compete with them for inorganic nutrients (Joint et al. 2002). In many diatom-bacteria associations, it is commonly observed a commensal-competitive continuum (Amin, Parker, and Armbrust 2012). The diatom-bacterium relationships described in this model system revealed a putative commensal phase mainly in the last part of the cultivation, in which bacteria seem to benefit from the extracellular organic products released by *P. tricornutum*, increasing their cell number.

### Population dynamics in cell-free spent medium

As it has been previously shown that diatoms can actually benefit from bacterial secreted compounds (Croft *et al*. 2005), we next asked whether the absence of a positive effect of bacterial cells on *P. tricornutum* could be due to some higher-order interactions in the co-culture that hides the potential beneficial effects of bacterial presences. Hence, to study the potential role of secreted bacterial substances, we performed diatom growth experiments in the presence of two different amounts of cell-free supernatants obtained from bacterial cultures (spent medium). *P. tricornutum* was cultured in 50% and 100% of spent bacterial medium. *P. tricornutum* grown in 50% diluted spent bacterial medium showed a rapid increase in cell number with respect to the diatom grown in non-diluted spent bacterial medium (100%) as well as in fresh Schatz salts (Papa et al. 2007) medium with some modifications (hereafter SS medium), the diatom positive control (Supplementary Figure S2). Having shown that the 50% diluted bacterial spent medium can boost the algal growth in the exponential phase (7-14 days), we can infer that *P. tricornutum* was able to use some soluble substances or metabolites present in the bacterial cell-free supernatant, in addition to all the macronutrients contained in the remaining half of synthetic medium.

These results were confirmed by the assessment of chlorophyll *a* content (Figure 1B). Indeed, also in this case, the chlorophyll *a* produced by the diatom grown in 50% diluted spent bacterial medium was higher than controls. These findings indicate that *P. haloplanktis* has the potential to promote *P. tricornutum* growth, but this does not happen when the two microorganisms are cultured together. Bruckner *et al*. (2011) showed that different amounts of cell-free bacterial supernatant were able to increase the growth of *P. tricornutum*, depending on the concentration of bacterial spent medium, suggesting that soluble factors released from bacteria can control diatom growth.

We then aimed to confirm that *P. haloplanktis* can thrive on *P. tricornutum*-derived compounds either in the form of dead *P. tricornutum* cells or released photosynthates. *P. haloplanktis* was firstly cultivated inside two different percentages of spent *P. tricornutum* medium (50% and 100%). *P. haloplanktis* grown in 50% diluted spent diatom medium showed increased growth in comparison to the bacterial negative control (Supplementary Figure S3A). Conversely, we observed that *P. haloplanktis* cultivated in 100% of spent medium exhibited a drastic reduction in the growth starting from the early stages of cultivation. We can presume that i) the spent diatom medium may contain extracellular photosynthates necessary for bacterial growth (observed with 50% of spent medium), but a higher percentage (such as 100%) could accumulate some substances that inhibit the growth of the bacterium or ii) algal growth may have sequestered the mineral nutrients necessary for bacterial subsistence.

Amin *et al*. (2012) argue that “diatoms seem able to “cultivate” their phycosphere by releasing organic-rich substances utilized by some bacteria”. Dissolved organic compounds (DOC) represent some of the main substrates provided by autotrophic diatoms to heterotrophic bacteria, and the diversity of these algal exudates likely play an important role, as a selective force, in shaping a diverse associated bacterial community. Recent research has highlighted how the same algal products could have opposite effects on different bacterial groups in order to select and modulate the bacteria associated with the diatoms. Diatom exudates include both central metabolites, accessible to the majority of bacteria, and also specific secondary metabolites able to promote the growth of selected bacteria and disadvantage others (Shibl et al. 2020).

Afterwards, we checked the capability of the bacterium to thrive on dead diatom cells. Accordingly, *P. haloplanktis* was successfully grown inside a medium containing diatom-autoclaved biomass as a substrate (Supplementary Figure S3B), revealing that the dead biomass of *P. tricornutum* was an excellent carbon source substrate. This result confirms that also dead phytoplankton cells are a source of organic matter for bacteria, which colonize such material to form a detritosphere (Richardson and Jackson 2007), and finally, thanks to enzymatic hydrolysis, convert detritus to DOM. Therefore, the freshly lysed diatoms represent one organic matter hot point that can sustain a vigorous growth of bacteria (Farooq Azam and Malfatti 2007).

From this whole body of data, we concluded that *P. tricornutum* has the potential to readily and efficiently sustain bacterial growth in the tested conditions, despite it is not yet possible to understand whether this is due to released photosynthates or the *P. tricornutum* dead biomass (or both). Conversely, *P. haloplanktis* did not affect the growth of *P. tricornutum* when these two microorganisms were grown together, although a positive effect of *P. haloplanktis* on the diatom was observed when the latter was grown on a (diluted) spent medium of the bacterial pure culture.

### A model for the *P. tricornutum*-*P. haloplanktis* co-culture

The growth data described above allowed us to infer a diatom-bacterium interaction network (Figure 2A). According to this network, *P. tricornutum* growth is made possible by light and CO_2_. The diatom sustains the growth of the bacterium by providing the required carbon sources (DOM, Figure 2A). Based on experimental observations, we postulated that the diatom has two alternative (but not exclusive) ways to provide nutrients to *P. haloplanktis*, either in the form of dead biomass (*DOM*_*B*_ or in the form of photosynthetic exudates (DOM_E_). In both cases, these nutrients can be taken up by bacterial cells and sustain their growth. We also included the possibility that part of the DOM pool (*DOM*_*B*_ + DOM_E_) is not taken up by the bacteria and leaves the system (empty set symbols in Figure 2A). Finally, we included a death rate for both *P. tricornutum* and *P. haloplanktis*.

**Figure 2:**
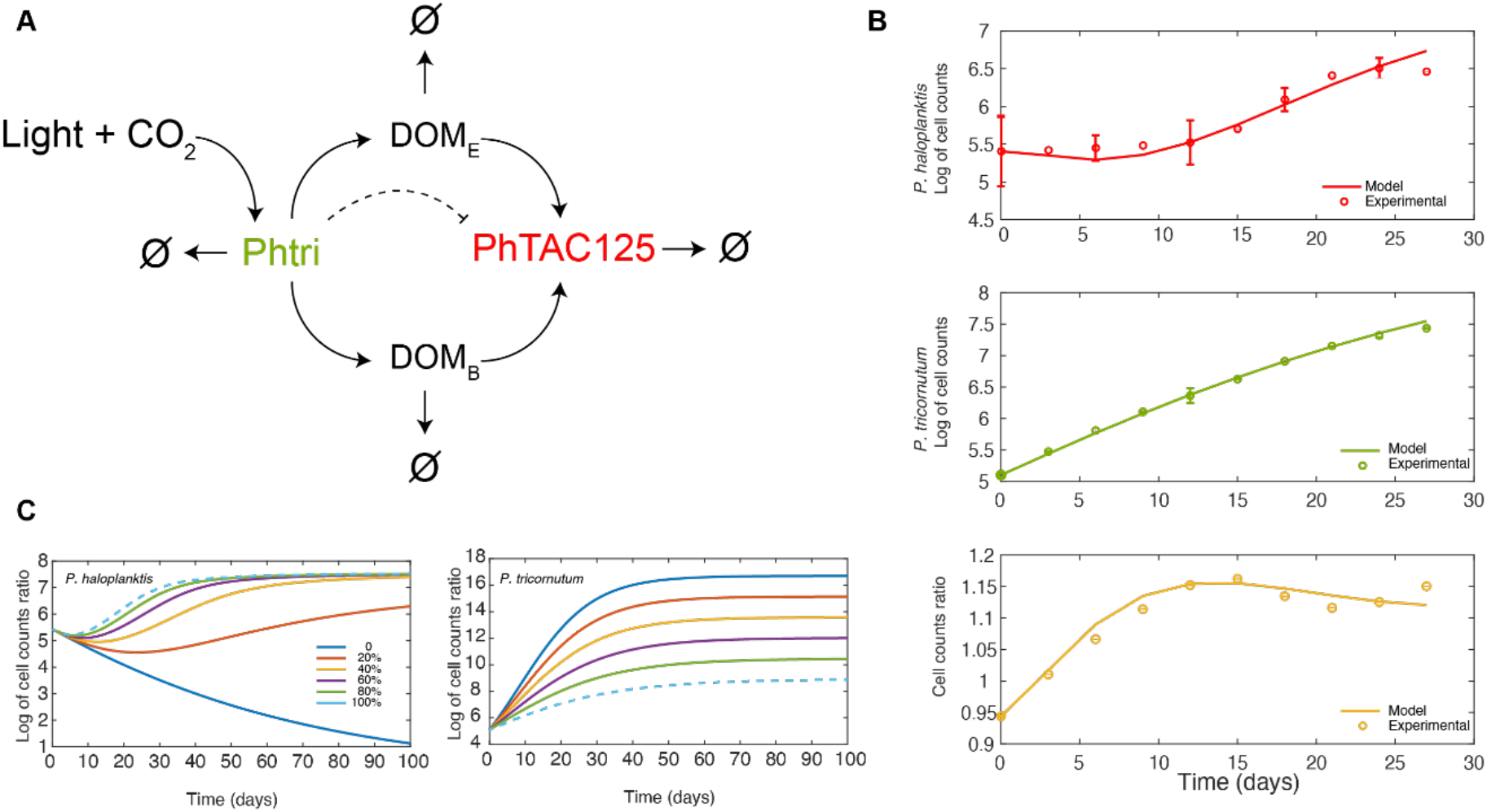
A) Hypothetical network explaining the co-culture interactions and dynamics. Phtri and PhTAC125 are abbreviations for *P. tricornutum* and *P. haloplanktis*, respectively. B) Simulation outcomes (continuous lines) and comparison with experimental (interpolated) data (empty circles). C) Simulations of bacterial and diatom growth with different exudates production rates (λ) of *P. tricornutum*, from 0 to 100% of the original fitted value. In this case, microbial growth was simulated for a period of 100 days to allow all the species to geta as close as possible to their steady state. The dashed lines represent the simulation with the original, estimated parameters.

The model is accounted for by the following set of ordinary differential equations, describing the change in time of the concentration of each species. In this formulation of the model, D and B respectively represent diatom and bacterial cell concentration, whereas light and CO_2_ were considered unlimited and thus not included in the actual equations implementing the network.

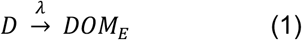

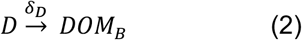

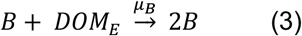

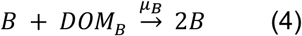

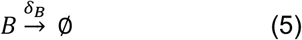

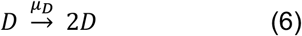

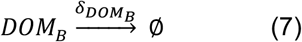

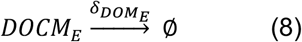

From this set of chemical equations, we derived the following set of ordinary differential equations:

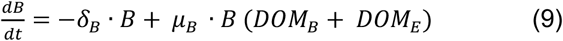

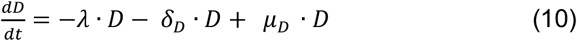

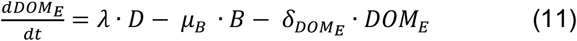

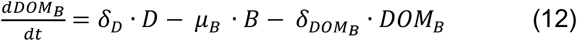

This model is based on a generalized logistic (Verhulst) growth model (Vogels et al. 1975) as already done by Moejes et al. (2017) to model the dynamics of a culture embedding *P. tricornutum* and a complex microbial community. Additionally, as *P. haloplanktis* is feeding on DOM released by *P. tricornutum* as the only carbon source, we included a Monod-type kinetic (Monod 1949) to reflect this dependency. Accordingly, the rates of Eq. 9-12 can be formalized as:

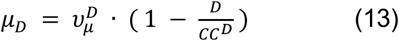

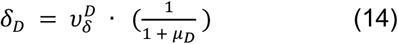

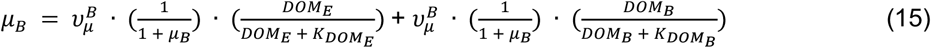

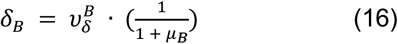

Where *CC* represents carrying capacities for the diatom (*CC*^*D*^) and bacterium (*CC*^*B*^), v the maximal rates for each growth or nutrients consumption processes, *DOM*_*E*_ and *DOM*_*B*_ the concentration of dead and exudated nutrients, respectively, and 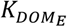 and 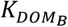 their corresponding Monod coefficients regulating nutrients exploitation. The rate at which diatoms (and phytoplankton in general) release organic carbon in the surrounding environment is the subject of an intense debate. Models have been proposed in which this rate (λ in our case) is either i) kept constant throughout the growth or ii) expressed as a function of the cellular state during the experiment. Model fitting on experimental data showed that both approaches could accurately describe the dynamics of the growth (Omta et al. 2020). Here we have interpreted the almost linear growth curve of *P. tricornutum* (Figure 1A and 2B) as a sign of an overall healthy population and thus decided to maintain a constant rate of *DOM*_*E*_ throughout the simulation. The model described here was fitted to the experimental data obtained from the co-culture experiments (Supplementary Table S2 for the details on model parameters). To increase the resolution power of the model and to ensure an efficient data fitting, we used an interpolation function (see Material and Methods) on the measured values of cell counts to increase the available points.

As shown in Figure 2B, the model fits well with the experimental data for bacterial, diatom and cell ratio experimental time points. The goodness-of-fit was computed for the two species (D and B) and their cell count ratio using the coefficient of determination (R^2^). We obtained an R^2^ of 0.977 (p-value = 1.104e-06), 0.99 (p-value = 2.324e-12) and 0.97 (p-value = 2.995e-06) for the fit to bacterial, diatom and cell count ratio, respectively. This analysis revealed a satisfactory precision of the model in describing the actual interactions between *P. tricornutum* and *P. haloplanktis* in the co-culture.

As seen before, both released exudates and diatom dead biomass can efficiently sustain bacterial growth. Having distinguished between exudate- and dead biomass-derived DOM (*DOM*_*E*_ and *DOM*_*B*_, respectively) gave us the possibility to evaluate the role of these two carbon sources in the predicted interaction network. Our model predicts that the growth of the bacterium relies almost entirely on actively produced *DOM* by the diatom (*DOM*_*E*_) with a very marginal role played by *DOM*_*B*_. This is further confirmed by the results of the simulations shown in Figure 2C, where we varied the rate of photosynthates release (λ) from 100% of the predicted value (obtained through data-fitting in the original simulation) to the 0% of the same value. Lower λ values negatively influence the growth of *P. haloplanktis*, especially for what concerns the initial lag time and increasing the discrepancy between simulated and experimental data. Running the same simulation on *δ*_D_ (the rate of *DOM*_*B*_ production that is coupled to *P. tricornutum* death rate) displayed no effect on the growth of the bacterium (data not shown).

Another main feature of the co-culture experiment is the remarkably long lag time of *P. haloplanktis* as 14 days are necessary to see a significant increase in bacterial cell count. This lag time may be due to either the necessity of *DOM* accumulation in the environment before the bacterium is actually capable of using it or a negative influence of *P. tricurnutum* on bacterial growth in the initial stages of the experiment (dashed line in Figure 2A). This latter scenario has been previously modeled to account for microbial inhibition in a two-or three-player co-culture (Cornu et al. 2011; Costa et al. 2019; Mickalide and Kuehn 2019).

Thus, we introduced the hypothesis according to which *P. tricornutum* exerts a negative control over *P. haloplanktis* growth in our model. Specifically, following the approach used in (Mickalide and Kuehn 2019) we modified Eq. 9 and as follows:

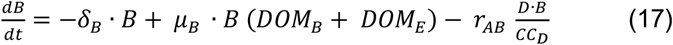

Here, the last term on the right-hand side of the equation takes into consideration the eventual inhibition of the diatom over *P. haloplanktis* according to an inhibition constant *r*_AB_ that was estimated through data fitting (starting from an initial guess of 0.3 reported in (Mickalide and Kuehn 2019)). This modified version of the model gave an overall goodness-of-fit (R^2^) of 0.61 (p-value = 0.05) and 0.91 (p-value = 0.0002) for bacterial and diatom growth, respectively (Supplementary Figure S4). A goodness-of-fit resembling the one observed for the original model (Figure 2B) was obtained for very low values of *r*_*AB*_ (*r*_*AB*_ = 9*10^−3^) that basically override the weight of the newly introduced term. Hence, according to these data, the inhibition of the diatom on the growth of the bacterium does not explain the initial growth lag of *P. haloplanktis*, suggesting that the time required for nutrient accumulation is a much more likely explanation for this co-culture.

## Conclusions

The complexity of microbial assemblages in aquatic ecosystems is due to the vast array of possible interactions among the players of the so-called microbial loop. This hampers the elucidation of the intimate functioning of the associations occurring between phototrophs and heterotrophs. In this work we have reduced the microbial loop to its lowest terms, i.e. a controlled co-culture of a phototrophic eukaryote and a chemoheterotrophic bacterium. By doing so we were able to account for the role of each of the two players in the established co-culture and, in particular, to suggest a flow of carbon sources from the diatom to the bacterium. This was further confirmed when the supposed network of interactions was implemented into a mathematical model that satisfactorily explained the dynamics of the co-culture. The mathematical model described in this work is rather simple compared, for example, to the one presented in Moejes et al. (2017). However, it represents a solid (i.e. guided by experimental growth data) basis for future upgrades, for example the inclusion of other microbes in the community, or of environmental perturbations (e.g. ocean acidification) to study their effects on the physiology of the co-culture. Finally, the fact that this synthetic community was built using model organisms for which a large body of knowledge already exists (including complete genome sequences, tools for their genetic manipulation, genome-scale metabolic reconstructions, mutant libraries), will permit to take the approach described in this work even further, assessing, for example, the role of specific genes and/or specific nutrients in the context of this microbial association.

## Material and methods

### Strains used and (co-)culturing methodologies

The axenic culture of the microalga *P. tricornutum* strain CCAP 1055/15 was purchased from the Culture Collection of Algae and Protozoa (CCAP, Scotland, United Kingdom) and cultured in f/2 medium with vitamin B_12_ (Guillard and Ryther 1962) at 20 °C under continuous illumination (15 µmol photons m^-2^ s^-1^) in static conditions inside an artificial climate incubator without CO_2_ supplementation. Axenic cultures were verified by microscopy observation and by inoculating samples in Marine Broth (MB) (Condalab, Spain), in a saline solution containing glucose (10 g/l) and in Luria Bertani (LB) (Malke 1993) broth with increased amount of NaCl (30 g/l) for 72 h at 27 °C in dark conditions.

Axenic algal cultures were periodically checked for the presence of bacteria by microscopy observation and plating on Marine Agar (MA) (Condalab, Spain). *P. tricornutum* cultures were maintained in our lab by transferring 3% of the culture volume to fresh medium every 4 weeks.

The antarctic bacterial strain *P. haloplanktis* TAC125 was obtained from the Institute Pasteur collection (CIP, Paris, France). This model bacterium was typically grown in MA plates or in Marine Broth (MB) (Condalab, Spain) incubated at 20°C under aerobic conditions.

Since, for the first time, these two model organisms were to be grown together, it was established a culture medium that would enable the growth of both model microorganisms to grow as single and co-cultures.

After in-depth preliminary tests (data not shown), for growth experiments were selected Schatz salts (SS) medium (Papa *et al*. 2007) with some modifications: 1 g/l KH_2_PO_4_, 1g/l NaNO_3_, 20 g/l sea salts, 0.2 g/l MgSO_4_ × 7H_2_O, 0.01 g/l FeSO_4_ × 7H_2_O, 0.01 g/l CaCl_2_ × 2H_2_O, 1ml/l trace elements stock solution (for 1 L of trace elements stock solution: 4,16 g Na_2_EDTA, 3,15 g FeCl_3_ × 6H_2_O, 0,01 g CuSO_4_ × 5H_2_O, 0,022 g ZnSO_4_ × 7H_2_O, 0,01 g CoCl_2_.6H_2_O, 0,18 g MnCl_2_ × 4H_2_O, 0,006 g Na_2_MoO_4_ × 2H_2_O) and adjusted to the pH of 7.

### *Pseudoalteromonas haloplanktis* TAC125 - *P. tricornutum* co-culture

The preculture of *P. haloplanktis* for co-cultivation experiments was grown for 3 days, in SS medium supplemented with L-glutamic acid (11 g/l) as the only C source in a 100 ml flask with a working volume of 25 ml, incubated at 20°C in the dark, with shaking at 100 rpm.

The co-culture was obtained by adding the bacterial preculture, washed twice by centrifugation at 4000 rpm for 4 min, up to a final density of 10^5^ cell/ml to the fresh culture of *P. tricornutum* prepared at the density of 2 × 10^5^ cell/ml in SS medium with no additional carbon source. Diatoms were inoculated from a growing stock culture of the axenic *P. tricornutum* in f/2 with vitamin B_12_.

In addition to the bacterial-diatom co-cultures, control cultures were prepared: *P. tricornutum* alone in SS medium as diatom control; *P. haloplanktis* alone in SS medium without C source and *P. haloplanktis* alone in SS medium containing additionally L-glutamic acid (11 g/l) as the only carbon source, as bacterial negative and positive control respectively. The co-cultures and all control cultures were grown in triplicate (final volume of 50 ml) in 100 ml flasks, incubated at 20°C under continuous illumination 15 µmol photons m^-2^ s^-1^, with shaking at 100 rpm. This co-culture experiment was conducted for 28 days.

### Growth experiments in cell-free spent medium

Additionally, two different growth experiments were performed: the diatom *P. tricornutum* was cultivated inside the bacterial spent medium, while the bacterium *P. haloplanktis* was grown inside the diatom spent medium. The bacteria were grown in SS medium with L-glutamic acid and incubated for 10 days in 1 L flask. Then, the 10-days-old bacterial culture was centrifuged, and the supernatant (spent bacterial medium) was sterilized in autoclave, adjusted to pH 7 in sterile conditions, and used to set up the growth experiment.

Diatoms were cultivated inside two different percentages of spent bacterial medium: 100% of non-diluted spent medium and 50% (v/v) of diluted spent medium (in fresh SS medium). Diatoms cultures in SS fresh medium were used as controls. The diatoms were inoculated at the cell density of 4 × 10^6^ cell/ml.

The same methodology was used for the cultivation of *P. haloplanktis* inside the diatom spent medium, obtained from a 20-days-old diatom culture in SS medium. Bacterial cultures were set up as negative and positive controls (as described above). All the cultures of the growth experiments in cell-free spent media were cultivated under the same light, temperature and shaking conditions described above.

The bacterium was also grown in the presence of dead-autoclaved biomass of *P*.*tricornutum* as the only carbon source for 28 days. *P. tricornutum* culture (dry weight: 1.66 g/l) was centrifuged, the biomass was autoclaved and then added to SS medium.

### Growth measurements

Growth of microalgae and bacteria under all conditions (co-culture and growth experiments in spent media) was monitored regularly every 7 days (from day 0 to day 28, 5 time points).

Diatom growth was determined by microscopic cell counts, using Thoma haemocytometer and by measuring chlorophyll *a* content following (Chen et al. 2011) method. Bacterial growth was measured by counting colony-forming units on MA plates.

### Model development and fitting

The model describing the bacterium-diatom co-culture was developed in MATLAB and the scripts are available at https://github.com/combogenomics/MicrobialLoop. To increase the set of points for data fitting, we used the *interp1* interpolation function implemented in MATLAB on the measured values of cell counts. The deterministic system was simulated by numerically integrating differential equations using the Matlab built-in (2019a) function ode45. To estimate the unknown parameters of the model from experimental data we used a stochastic curve-fitting *in-house* Matlab software. The algorithm is based on the paper by Cardoso et al. (Cardoso, Salcedo, and Feyo de Azevedo 1996) and consists in the combination of the non-linear simplex and the simulated annealing approach to minimize the squared deviation function.

## Notes

https://github.com/combogenomics/MicrobialLoop

### Competing Interest Statement

The authors have declared no competing interest.

