## Supplementary Figure S1 for "Scaling down the microbial loop: data-driven modelling of growth interactions in a diatom-bacterium co-culture"

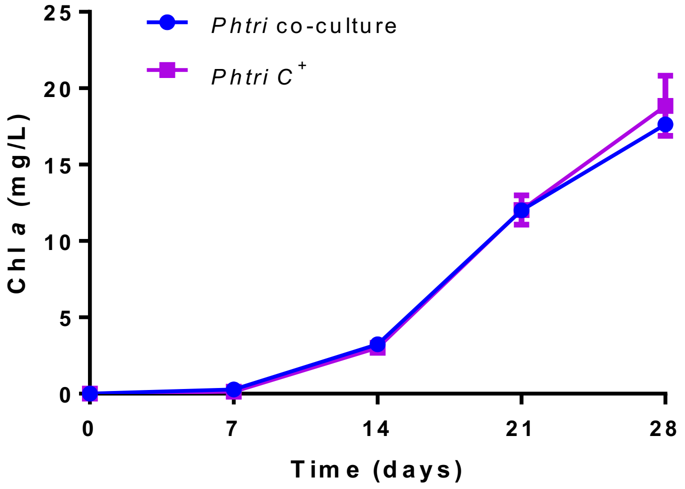


**Figure S1:** Chlorophyll *a* content in *Phaeodactylum tricornutum* (*Phtri*) grown in co-culture with the bacterium *Pseudoalteromonas haloplanktis* TAC125 and in the control. Co-culture and *P. tricornutum* positive control were cultured in SS medium. Error bars, standard deviation of three biological replicates.

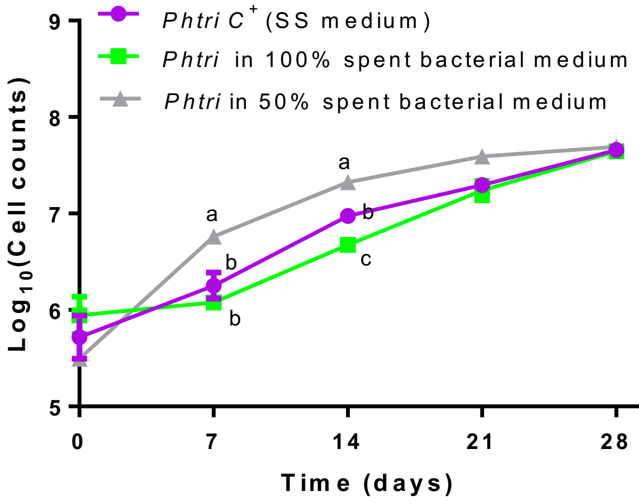


**Figure S2:** Growth curves of the diatom *P. tricornutum* (*Phtri*) in spent bacterial medium (grey, 50% diluted bacterial spent medium, green 100% bacterial spent medium, violet, control grown in fresh SS medium). Error bars, SEM (Standard Error of the Mean) of duplicate cultures. Different letters describe significant difference (ANOVA, Post-test: Tukey's multi comparative test, $p<0.05$).

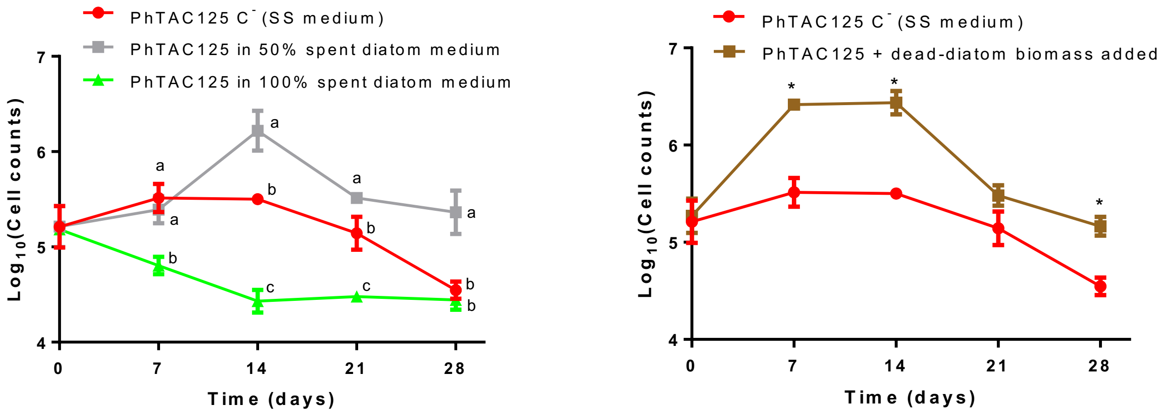


**Figure S3: A)** Growth curves of the bacterium *P.haloplanktis* TAC125 (PhTAC125) in 50% diluted and 100% of spent diatom medium. *P. haloplanktis* TAC125 negative control grown in SS medium. Error bars, standard deviation of triplicate cultures. Different letters describe significant differences (ANOVA, Post-test: Tukey's multi comparative test, $p<0.05$). **B)** Growth curves of the bacterium PhTAC125 grown in a medium containing diatom-autoclaved biomass. Error bars, standard deviation of triplicate cultures. The asterisk indicates significant difference (t-test, $p < 0.05$).

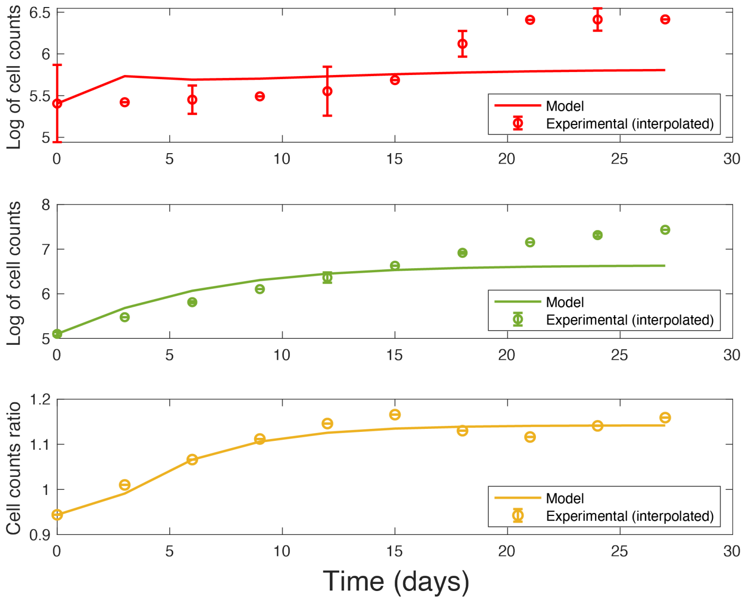


**Figure S4:** Simulation outcomes (continuous lines) and comparison with experimental (interpolated) data (empty circles) of a model implementing an inhibitory effect of *P. tricornutum* over *P. haloplanktis*. Colour codes as in Figure 2.

**Table S1**: Cell counts of *P. haloplanktis* TAC125 positive control in the co-culture experiment. Mean and standard error of three biological replicates.

| **Time (days)** | **Cell counts: CFU/ml** | |
|---|---|---|
| | **Mean** | **Standard error** |
| 0 | $1.26 \times 10^5$ | $5.45 \times 10^4$ |
| 14 | $6.7 \times 10^7$ | $1.82 \times 10^7$ |
| 21 | $8.28 \times 10^6$ | $1.13 \times 10^6$ |
| 28 | $2.92 \times 10^6$ | $6.97 \times 10^5$ |

**Table S2:** List of model parameters used in the model.

| **Parameter** | **Description** | **Fitted value** |
|---|---|---|
| $\upsilon_\mu^D$ | Maximal diatom growth rate | 0.0984 |
| $CC^D$ | Diatom carrying capacity | 16.7011 |
| $\upsilon_\delta^D$ | Maximal diatom death rate | 0.0000 |
| $\upsilon_\mu^B$ | Maximal bacterial growth rate | 0.1801 |
| $CC^B$ | Bacterial carrying capacity | 7.4410 |
| $K_{DOM_E}$ | Monod-like coefficient for $DOM_E$ | 10.8307 |
| $\upsilon_\delta^B$ | Maximal bacterial death rate | 0.0178 |
| $\lambda$ | $DOM_E$ release rate | 0.0459 |
| $K_{DOM_B}$ | Monod-like coefficient for $DOM_B$ | 0.2400 |
| $\delta_{DOM_B}$ | $DOM_B$ death rate | 0.0000 |
| $\delta_{DOM_B}$ | $DOM_E$ death rate | 0.0000 |
| $r_{AB}$ | Diatom-bacterium inhibition constant | 0.0010 |
